# Highly correlated activity across higher-order thalamic nuclei in awake and anesthetized states

**DOI:** 10.1101/2025.10.03.680235

**Authors:** Toon Brouwer, Mathis Bassler, Cyriel M.A. Pennartz, Mototaka Suzuki

## Abstract

The thalamus is an egg-shaped structure deep inside the brain and is of central importance for our cognitive functions. The thalamus can be divided into first- and higher-order (HO) nuclei. Unlike first-order thalamic regions which are known to primarily relay sensory signals to the neocortex, the functions of HO thalamic nuclei remain far less clear. Although previous studies indicate HO thalamic nuclei influence cortical processing and consciousness, most studies examined single thalamic nuclei in isolation. To investigate the relation between distinct HO thalamic nuclei, we recorded local field potentials (LFP) and multi-unit activity (MUA) across multiple thalamic nuclei simultaneously in awake and anesthetized mice with sensory stimulation. The effect of voluntary locomotion was also studied in the awake state. Surprisingly, we found that both LFP and MUA were strongly correlated across HO nuclei in both states, which cannot be explained by volume conduction alone but suggests the existence of shared synaptic inputs. Furthermore, in the awake state locomotion and sensory stimuli activated many HO nuclei, including those that have not classically been regarded as sensory or motor. These findings challenge the classical view of the thalamus being the aggregate of isolated, independent nuclei and suggest that the thalamus serves as an ideal bottleneck whose coordination resulted in effective orchestration of the entire brain.

## Introduction

The thalamus is an important brain structure that is shared by all vertebrate species and consists of distinct nuclei^1,2^. Thalamic nuclei can anatomically be divided into first-order and higher-order nuclei^1^. First-order thalamic regions are known to primarily relay sensory signals from sensory peripherals to the neocortex^1^. Higher-order (HO) thalamic nuclei, in contrast, predominantly receive “driving” or “primary” inputs from the cerebral cortex^1,2^. Apart from the anatomical feature, however, the functions of HO thalamic nuclei are much less known, compared to the first-order nuclei. Different HO nuclei are involved in a wide range of distinct circuits related to a multitude of functions. For example, the posterior medial nucleus (POm) has been linked to sensory perception and perceptual discrimination^3,4^; the pulvinar to prediction error signaling^5,6^; the mediodorsal nucleus (MD) to executive control^7^; and the ventromedial nucleus (VM) to arousal and goal-directed action initiation^8,9^. These previous findings suggest that HO thalamic nuclei may support a wide variety of distinct, specialized functions.

Despite these functional differences, interestingly, many HO nuclei have been commonly associated with global brain states and consciousness. HO nuclei such as the POm, MD, pulvinar, as well as intralaminar nuclei such as the central lateral (CL), centromedian (CM), paraventricular (PVT), and parafascicular (PF) nucleus, have all been implicated in aspects of consciousness^10–17^. This functional overlap is striking given the considerable diversity among HO nuclei: they project to and receive inputs from a wide range of brain regions while lacking local excitatory recurrent connections^18^, which is in stark contrast with the cerebral cortex that has abundant local excitatory recurrent connections.

Despite the previous findings in HO thalamic nuclei, most previous studies examined single thalamic nuclei in isolation; therefore, the functional relationships between different HO nuclei across conscious states are unknown. To address this, we measured neuronal activity (LFP and MUA) in multiple thalamic nuclei (HO and other thalamic nuclei) simultaneously using a four-shank multi-channel electrode array *in viv*o. We measured responses to sensory stimuli presented during both awake and isoflurane-anesthetized states. We also measured thalamic activity when the awake animal was voluntarily locomoting or standing still. We surprisingly found that the neuronal activity (both LFP and MUA) of HO thalamic nuclei are strongly correlated, which cannot be explained by volume conduction alone and instead suggest the existence of shared inputs that coordinate the activity across HO nuclei. An emerging view is that the thalamus serves as an ideal bottleneck whose coordination results in effective orchestration of the entire brain. In Discussion, we will discuss the implications of our findings.

## Results

We recorded LFP and MUA in mice using a four-shank electrode array—with 8 electrodes per shank, that spanned 1,200 μm horizontally and 1,400 μm vertically (Fig. 1a)—enabling us to sample a broad range of thalamic nuclei simultaneously in every animal (Fig. 1b). Post-hoc histological analysis showed that about half (178/352) of our recordings were from the MD, ventrolateral (VL), laterodorsal (LD), anteroventral (AV), CL, anteromedial (AM), ventral posteromedial (VPM), thalamic nuclei (Fig. 1c), although VPM is a first-order (FO) nucleus. Twenty-six recordings were also from the lateral and medial habenula (LHB/MHB).

**Figure 1:**
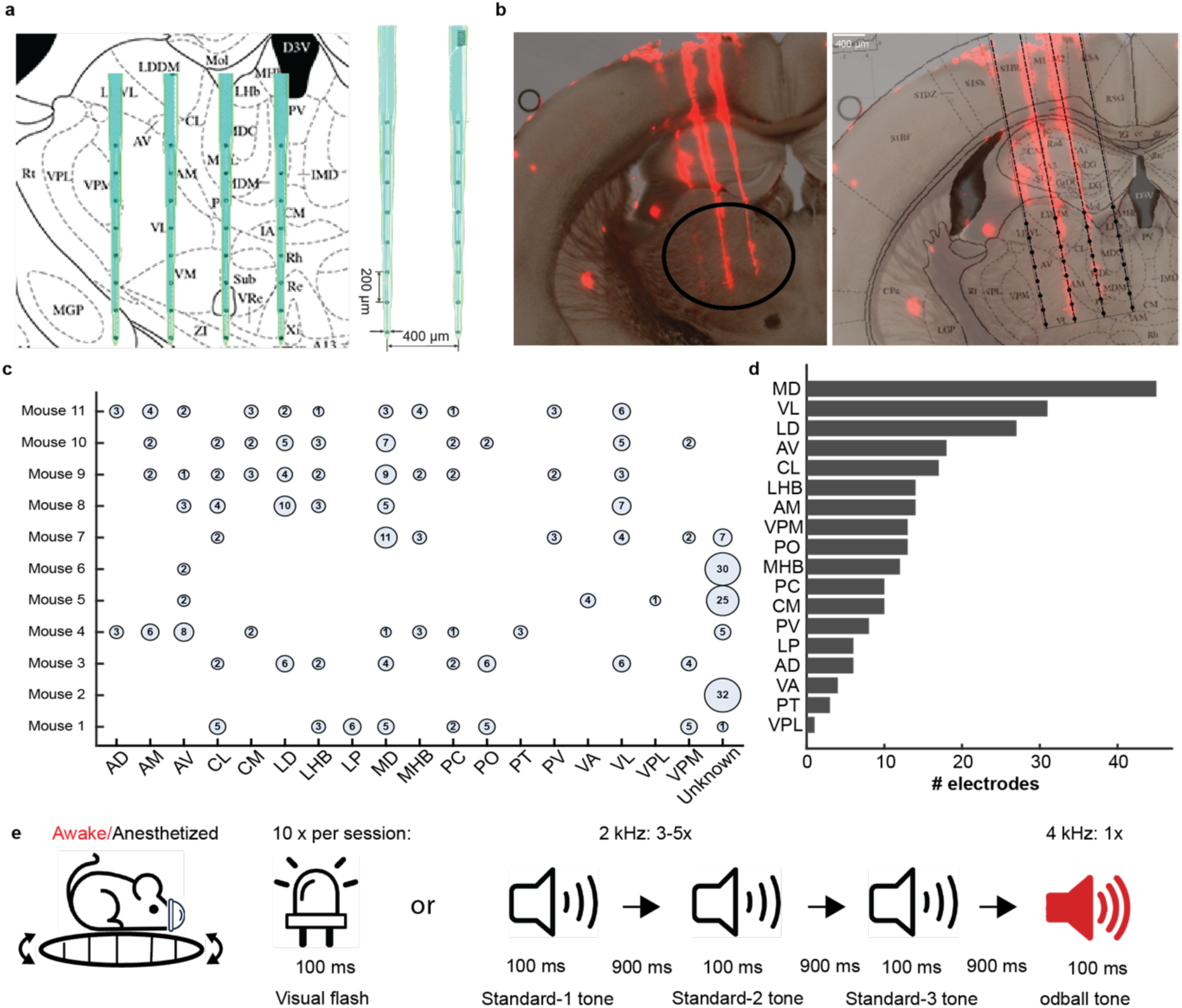
Simultaneous recording of multiple thalamic nuclei. (a) Left: Recordings used a four-shank probe with 200μm intervals between electrodes within each shank and 400μm inter-shank intervals. (b) An example histological image showing the target location. Right: An illustration of the target region of the probe, recording in many higher-order thalamic nuclei concurrently. (c) Summary of the recorded thalamic nuclei in 11 mice. The number and size of the circle indicate the number of recording sites within the nucleus indicated. Recording sites that did not clearly match the location of any thalamic nucleus were labelled as ’unknown’. (d) Summary of the total number of recording electrodes from each thalamic nucleus. (e) The experimental paradigm. Thalamic activity was recorded in both awake and anesthetized states. In each session animals were presented with 100 ms visual flashes of light or a random number of tones (3-5 standard 2 kHz tones followed by one oddball tone at 4 kHz with an inter stimulus interval (ISI) of 900 ms). Moreover, when awake, animals could walk or run on a treadmill at will.

In both anesthetized and awake states, mice were presented with visual stimuli (100ms single flashes of white LED light) and auditory stimuli consisting of an oddball paradigm with a random number (from three to five) of 2-kHz standard tones followed by one 4 kHz deviant (Fig. 1d). When awake, all mice were allowed to voluntarily locomote on a linear treadmill.

**Table 1:** Overview of Nuclei Abbreviations.

| Abbrev. | Full Name | Abbrev. | Full Name |
| --- | --- | --- | --- |
| **MD** | Mediodorsal nucleus | **VL** | Ventrolateral nucleus |
| **LD** | Lateral dorsal nucleus | **AV** | Anteroventral nucleus |
| **CL** | Central lateral nucleus | **LHB** | Lateral habenula |
| **AM** | Anteromedial nucleus | **VPM** | Ventral posteromedial nucleus |
| **PO** | Posterior nucleus | **MHB** | Medial habenula |
| **PC** | Paracentral nucleus | **CM** | Centromedian nucleus |
| **PV** | Paraventricular nucleus | **LP** | Lateral posterior nucleus |
| **AD** | Anterodorsal nucleus | **VA** | Ventral anterior nucleus |
| **PT** | Paratenial nucleus | **VPL** | Ventral posterolateral nucleus |

### General anesthesia decreases multi-unit activity in thalamic nuclei

To examine how anesthesia influences the activity of thalamic nuclei, we recorded and analyzed the MUA and LFP in both awake and anesthetized state (Fig 2a). We observed that the mean MUA across all recording sites significantly decreased during anesthesia (p<0.001, two-sided Wilcoxon signed-rank test, n=11 mice, Fig 2b), whereas the root-mean-square of the LFP amplitude was not significantly affected (p=0.206, two-sided Wilcoxon signed-rank test, n=11 mice, Fig 2c). Subsequently, we examined the power spectrum of the LFP in both states to assess whether the effect of anesthesia was specific to a certain frequency range (Fig 2d). We found that the power in lower frequency bands (<4 Hz) was similar between anesthesia and awake, whereas in all higher ranges (>4Hz) the power seemed lower in anesthesia than in wakefulness. To quantify this, we computed the ratio of the power in the delta range (< 4Hz) over the power in the rest of the spectrum (>4 Hz cut off at 30 Hz) and found that this ratio was higher during anesthesia compared to wakefulness (p<0.001, linear mixed effects model, n=352 sites, Fig 2e), suggesting relatively higher Delta range activity (0.5-4 Hz) during anesthesia. This is consistent with the previous studies (increase of slow-wave activity during anesthesia)^19–21^.

**Figure 2:**
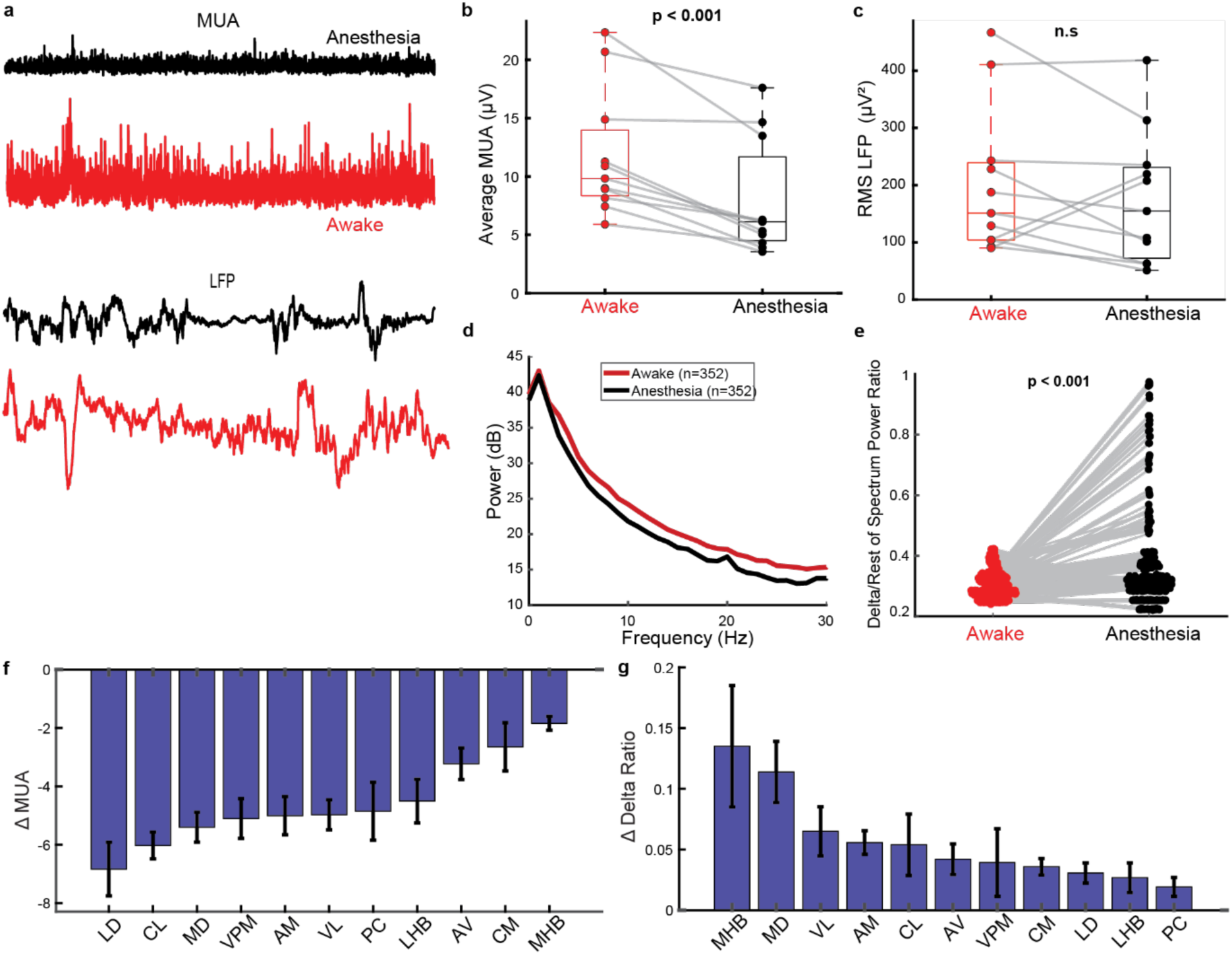
Anesthesia decreases the neuronal activity across HO thalamic nuclei. (a) Top: Example MUA trace of an electrode during anesthesia (black) and wakefulness (red). (b) MUA (average envelope amplitude) decreases during anesthesia (two-sided Wilcoxon signed-rank test, p<0.001). Each dot represents one animal (n=11) (average over time and recording sites). (c) LFP amplitude (root mean square, RMS) does not significantly decrease during anesthesia (p=0.206, two-sided Wilcoxon signed-rank test). Each dot represents one animal (n=11). Box whiskers indicate minimum and maximum values of the data. (d) Grand average power spectrum across all recording sites from all animals (n=352 sites). (e) LFP spectral profile changes during anesthesia. The plot shows the ratio of Delta (0.5-4 Hz) activity divided by the power in the rest of the sub-30 Hz spectrum to assess the relative prominence of low-frequency activity; each dot represents one recording site (n=352 sites). Anesthesia significantly increased the Delta ratio (linear mixed effects model, p<0.001). (f) Anesthesia decreases MUA across thalamic nuclei. The difference in average MUA (Anesthesia – Awake) is displayed for all nuclei that were recorded from at least 4 animals and from which at least 10 total recording sessions were available. Error bars indicate s.e.m. (g) Anesthesia increases the Delta ratio across thalamic nuclei. The difference in delta ratio (Anesthesia – Awake) is displayed for the same nuclei as in (f).

Next, we wondered whether these effects were consistent across the majority of thalamic nuclei or whether they were driven by only a few thalamic regions. The decrease in MUA was significant in all thalamic nuclei of sufficient data (>4 animals, >8 total recording sites) (all p<0.05, Wilcoxon signed rank test, FDR corrected), and the increase in the Delta ratio (Fig. 2g) was statistically significant in all thalamic nuclei (all p<0.05, Wilcoxon signed rank test, FDR corrected) but the first-order nucleus VPM.

### Thalamic activity is highly correlated during both anesthesia and wakefulness

While recording the HO thalamic LFP activity, we were surprised by the similarity of the signal measured at many electrodes—even the electrodes that are far distant (>1.2 mm) from each other (Fig 3a). We therefore calculated the correlation coefficient between the LFP of all electrode pairs in both anesthetized and awake states. The LFP strongly correlated in both states (wakefulness r=0.95, anesthesia r=0.97, both p<0.001, two-sided Wilcoxon signed-rank test against zero, n=11 mice, Fig 3b). Moreover, this strong correlation did not differ significantly between anesthesia and wakefulness (p=0.12, two-sided Wilcoxon-signed rank test against zero, n=11 mice, Fig 3b left). A possible reason for such a high correlation is volume conduction from regions inside or outside the thalamus. To examine this possibility, we performed multiple analyses. First, we used bipolar re-referencing (see Methods) to amplify the local signal and measured the correlation coefficient but now looked at the absolute value of the correlation coefficient (to prevent canceling out between positive and negative values introduced by bipolar re-referencing). In wakefulness, the mean bipolar-LFP correlation was r=0.30, and under anesthesia it was r=0.78 (both p<0.001, two-sided Wilcoxon signed-rank test, n=11 mice; Fig. 3b middle. In contrast to the monopolar LFP, the bipolar-LFP showed an increased amount of correlation between different electrodes in anesthesia, compared to wakefulness (p=0.02, two-sided Wilcoxon-signed rank test, n = 11 mice, Fig 3b middle). Lastly, we also measured the correlation coefficients of the MUA due to the localized nature of this signal and found similar amount of correlation between different electrodes (r=0.29 during wakefulness and r=0.30 during anesthesia, both p<0.001, two-sided Wilcoxon signed-rank test, n = 11 mice) but no significant difference in correlation between anesthesia and wakefulness (p=0.83, two-sided Wilcoxon signed-rank test, n = 11 mice).

**Figure 3:**
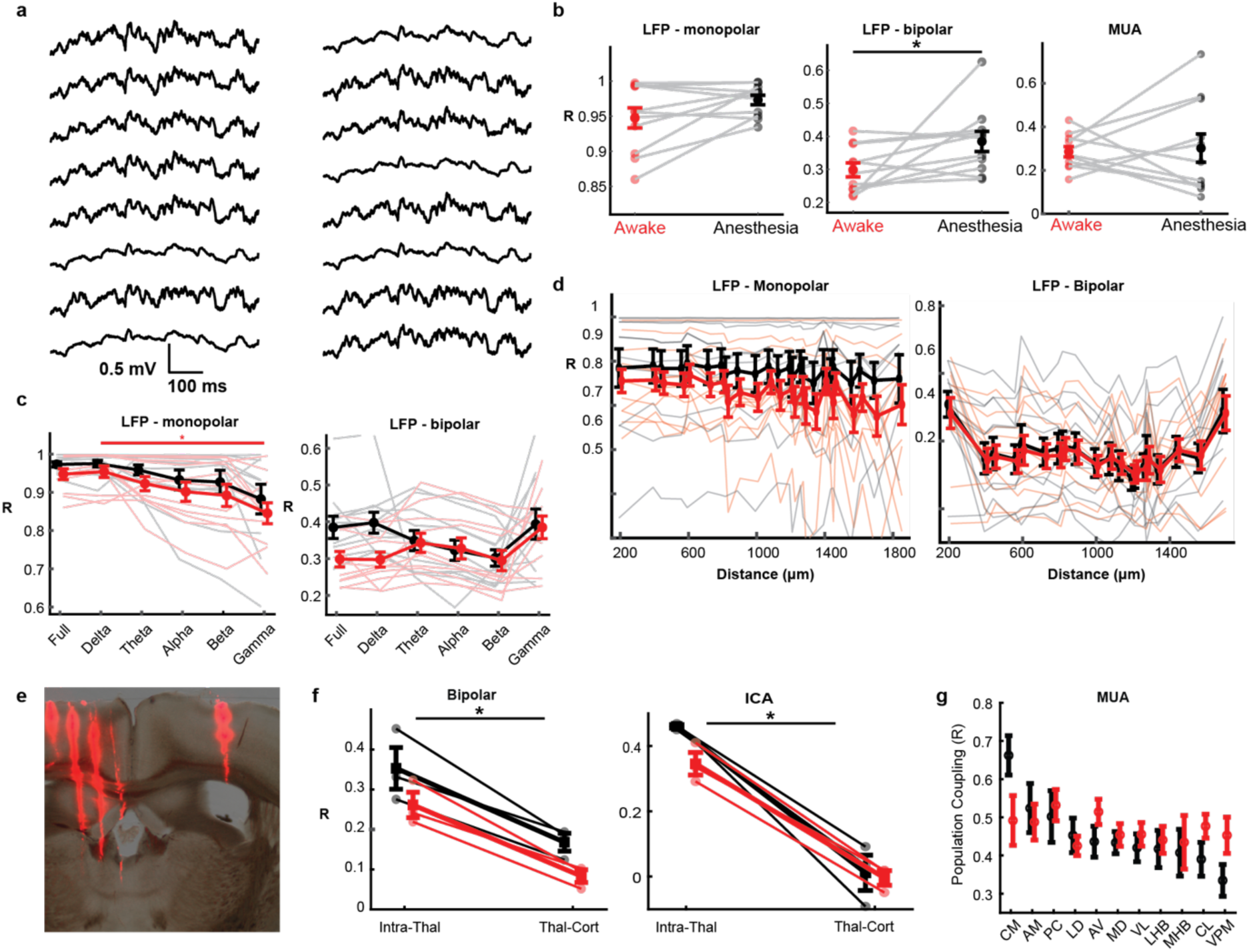
Highly correlated LFP across HO thalamic nuclei. (a) Example LFP traces recorded from electrodes across two shanks (distance: 400μm), showing the similarity between distant electrode sites. (b) Thalamic sites show highly correlated LFP. Left: Average correlation coefficients (R) between the monopolar-LFP signals averaged across all electrode combinations (n=11 mice). Error bars indicate s.e.m. for this and subsequent subfigures. Middle: Same for the bipolar-LFP. Right: Same for the MUA. (c) Correlation coefficients across different frequency bands. Left: average correlation across all electrode sites (n=11 mice) during wakefulness and anesthesia with the signal filtered in different frequency bands, showing no significant effect of frequency on correlation strength during anesthesia (p=0.39, Kruskal-Wallis test) but with a significant effect of frequency on correlation during wakefulness (p=0.021, Kruskal-Wallis test); the Gamma band (30-80 Hz) showed a lower correlation compared to the Delta band (p=0.021, Dunn-Sidak test). Right: same for the bipolar-LFP with no significant effect of frequency in both anesthesia and wakefulness (p=0.148 and p=0.186, Kruskal-Wallis test). (d) Distance dependency of the intra-thalamic correlation. Top: Correlation across absolute distance (Euclidean distance computed from vertical and horizontal distance) for both the monopolar- (left) and bipolar-LFP (right). The correlation remains high at increasing distances. Again the correlation remains high at increasing distances but does show a decrease with distance for the bipolar LFP (p<0.001 both states, β (slope) ≃-0.0002, linear mixed effects model) (right). (e) A photomicrograph of a brain section after recording HO thalamic nuclei with a 4-shank electrode array and a contralateral cortical area with a 1-shank linear 16-channel electrode array. (f) Intra-thalamic vs. thalamo-cortical correlations of bipolar LFP in awake and anesthetized states. (g) Significantly higher intra-thalamic correlations are maintained even after removing the globally correlated component calculated by Independent Component Analysis (ICA) applied to the pooled LFP data from HO thalamic nuclei and the contralateral cortical area shown in (e). (h) Recorded thalamic nuclei all have MUA signals that are highly correlated to the population average. Population coupling represents the average correlation between the MUA from a specific electrode site to the summed signal across all other sites. Error bars indicate s.e.m.

Next, we examined whether the strong LFP correlation between electrodes/sites was bound to a specific frequency band for both the monopolar- and bipolar-LFP. For the monopolar-LFP the correlation coefficient did not significantly differ between frequency bands during anesthesia (p=0.39, Kruskal-Wallis test, n=11 mice, Fig 3c left), but during wakefulness, there was a significant difference (p=0.021, Kruskal-Wallis test, Fig 3c left); specifically, the Gamma band showed a significantly lower average correlation between electrodes compared to the Delta band (p=0.021, Dunn-Sidak test, Fig 3c). In case of the bipolar-LFP the correlation coefficient did not significantly differ between frequency bands during both anesthesia and wakefulness (p=0.148 and p=0.186, Kruskal-Wallis test, Fig 3c right).

Next, we examined whether the LFP correlation depends on the distance between recording sites. Using a linear mixed effects model (see Methods), we found that the distance between electrodes along the shank (vertical distance) did not significantly determine their monopolar-LFP correlation during both anesthesia (p=0.1707, linear mixed effects model, Fig 3d left lower) and awake states (p=0.506, linear mixed effects model, Fig 3d left lower). Absolute distance—computed from both vertical and horizontal distances—was associated with a very marginal decrease in correlation in both states (β (slope) ≃ 10⁻⁶). This effect reached significance under both anesthesia (p=0.008) and wakefulness (p=0.004; Fig. 3d top left). In case of the bipolar-LFP correlation we found a significant (yet marginal) decrease of correlation with absolute distance in anesthesia (p<0.0001, β (slope) ≃-1*10⁻^5^), but not during wakefulness (p=0.201, linear mixed effects model, Fig 3d right upper). Thus, the distance between recording sites does not strongly affect the strength of correlation.

To examine whether the LFP correlation was specific to the thalamus, we also measured LFP activity in the cortex together with the thalamus (Fig 3e). We then compared the average correlation between LFP within the thalamus (intra-thalamic correlation) to the average correlation between the thalamus and the cortex (thalamo-cortical correlation). For the bipolar LFP, intra-thalamic LFPs were significantly higher than the thalamo-cortical correlation in both anesthesia (intra-thalamic correlation: r=0.33, thalamo-cortical correlation: r=0.17, t=4.56 p=0.045, two-sided paired t-test, n=3 mice) and wakefulness (intra-thalamic correlation: r=0.26, thalamo-cortical correlation: r=0.08, t=4.56 p=0.045, t=6.82 p=0.021, two-sided paired t-test, n=3 mice, Fig 3f left). As another attempt to control for volume conduction, we performed an independent component analysis (ICA) to find the component which was most closely related to the global average (across all cortical and thalamic recording sites) and subtracted this global component from all sites (see Methods for more details). Subsequently, we examined if the thalamic LFPs still showed significant correlations after removing the globally correlated component; indeed, we still found substantial intra-thalamic correlations during both anesthesia (r=0.41) and wakefulness (r=0.32) which were higher than the thalamo-cortical correlation in both anesthesia (t=4.83, p=0.04, two-sided paired t-test, n=3 mice, Fig 3g right) and wakefulness (t=13.06, p=0.006, two-sided paired t-test, n=3 mice, Fig 3g right). Two other analytical methods than ICA – Principal Component Analysis and common average re-referencing – also produced unique and similarly strong intra-thalamic correlation, compared to the thalamo-cortical correlation (Supplementary figure 1).

Additionally, we tested whether the correlations were specific to a particular group of thalamic nuclei only. We computed the population coupling (see Methods) to see how strongly the MUA in each electrode was correlated to the summed MUA across all recording sites (i.e., all simultaneously recorded thalamic nuclei). We found that MUA of recorded thalamic nuclei were all highly correlated to the population MUA (all >0.3, p<0.05, FDR corrected, Pearson Correlation tested against 0, Fig 3h), suggesting the coordinated activity between different thalamic nuclei.

### Sensory responses in thalamic nuclei are modality- and state-dependent

We next addressed the question of how thalamic nuclei respond to sensory stimuli in each conscious state as measured by MUA. Because of our finding that MUA power was reduced during anesthesia (Fig 2b), we expected reduced stimulus responses under anesthesia.

For each electrode, we calculated the mean MUA response for 500 ms after each stimulus onset as well as the activity when no stimulus was presented (Fig 4a & b, see also Methods). We then tested the significant responses recorded in each electrode in each state (Kruskal-Wallis test, see Methods). After correcting for multiple comparisons, we found 191/344 recording sites (from 10/11 mice) responsive in at least one state. We were surprised to find similar fractions of responsive sites in wakefulness (33%) and anesthesia (37%).

**Figure 4:**
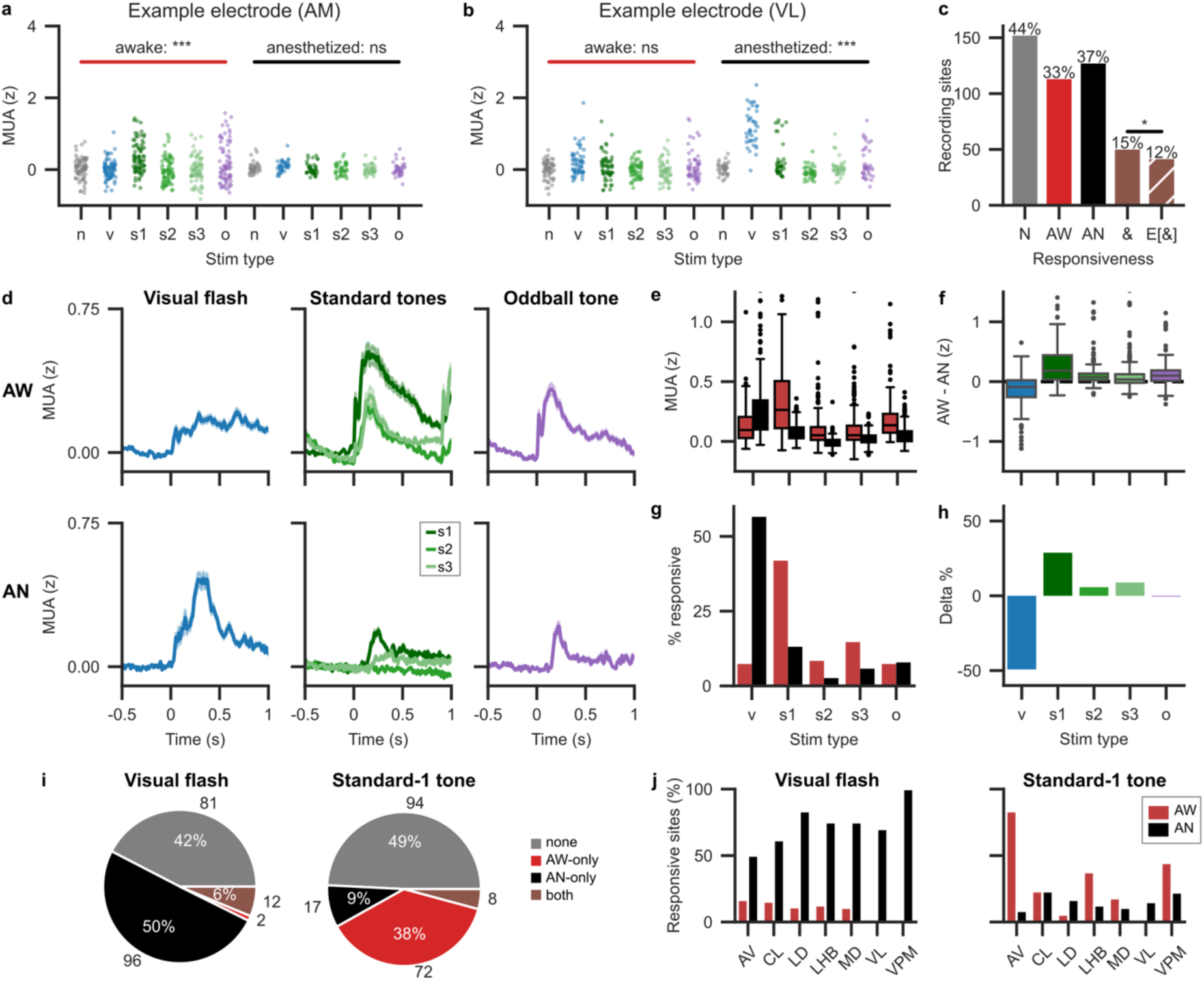
State-specific stimulus responsiveness across thalamic nuclei. (a) Example multi-unit activity (MUA) of an electrode in anteromedial nucleus (AM). This electrode shows significant responses only during wakefulness. Each dot represents the average z-scored MUA response in a single trial within 500 ms after stimulus onset. Colors indicate stimulus type. b (grey) = baseline, v (blue) = visual flash, s1 (dark green) = standard-1 tone, s2 (green) = standard-2 tone, s3 (light green) = standard-3 tone, o (purple) = oddball tone. *** = p < 0.001, Kruskal-Wallis test, FDR correction over 344 tests. ns = not significant. (b) Same as (a) but for an electrode in the ventral lateral nucleus (VL). Notice that the visually evoked activity is only during anesthesia. (c) Recording sessions of significant responses to stimuli as assessed by Kruskal-Wallis tests. Bar size indicates absolute number of recording sessions while percentages on top indicate the fraction of the total number of electrodes (n=344). N, not responsive in either state, AW, responsive (at least) during wakefulness, AN = responsive (at least) under anesthesia, & = responsive in both states, E[&], expected number of electrodes responsive in both states if probabilities of being responsive in one state were independent from probability of being responsive in the other state. The symbol ‘*’ indicates p < 0.05, Fisher’s exact test. Only 15% of electrodes were significantly responsive in both states which was only slightly above statistical independence. (d) Stimulus-triggered average z-scored MUA responses of all responsive sessions (n = 191) to different types of stimuli (columns) separated by state (row). Shaded area indicates s.e.m. (e) Boxplot of average MUA responses of all 191 responsive sites to all stimuli in both the anesthetized and awake state. Outliers (z>1.25) are not shown. Red: awake, black: anesthetized. Average MUA responses were significantly different between states for all stimuli (two-sided Wilcoxon rank sum test, all p < 0.001). (f) Boxplot of difference between awake and anesthetized MUA responses for each stimulus type of responsive sites. Outliers (z>1.25) are not shown. Awake MUA responses exceeded anesthetized MUA responses for all tones, but anesthetized MUA responses to the visual flash exceeded awake ones. (g) Fraction of stimulus-responsive sites as assessed by Mann-Whitney U test between stimulus and null responses. Colors are as in (e). (h) The fraction of responsive sessions in awake vs anesthetized states for each stimulus type. (i) Pie charts indicating stimulus responsiveness by state for visual and standard 1 stimuli. Most responsive sites were only responsive in one of the two states. Numbers outside of the circle indicate absolute numbers of responsive sessions. Abbreviations as in c. (j) Fractions of stimulus-responsive sites in anesthesia vs. awake states for six thalamic nuclei as well as the LHB for visual flash and standard-1 tone stimuli.

Interestingly, of the 191 responsive sites, the majority (140 sites) was only responsive during either wakefulness or anesthesia; only 15% (51/344) of sites were responsive in both states (Fig 4c). The observed fraction of sites that were responsive in both states was only modestly greater than the fraction predicted (12%) if responsiveness was totally independent across states (p = 0.028, one-sided Fisher’s exact test). These data suggest that the thalamic response strongly depends on the conscious state, and the response in one state does not guarantee the responsiveness in the other state.

We next looked at responses to five specific stimulus types: the visual flash, the first three repeated tones (called standard-1, standard-2, and standard-3 tone, respectively) and the subsequent oddball tone. We limited this analysis to the 191 sites that were significantly responsive in at least one state. First, we calculated the MUA response pooled across all recording sites to each stimulus type in each state (Figure 4d). We found significant differences of responses to all stimulus types between anesthesia and wakefulness (p < 0.001, two-sided Wilcoxon rank-sum test, n = 191 sites; Figure 4e). We found stronger auditory responses during wakefulness compared to anesthesia (Figure 4f). Surprisingly, the opposite was the case for the visual stimulus: light flashes evoked a much stronger response in thalamic nuclei during anesthesia than during wakefulness (Figure 4g&h).

Subsequently, we found the responsiveness of individual sites to each stimulus type (two-sided Mann-Whitney U test, α=0.05, see Methods). We found that only the visual stimulus under anesthesia (57% = 108/191 sites) and standard-1 tone during wakefulness (42% = 80/191 sites) evoked significant responses (less than 15% sites responded to other stimuli). We therefore focused our attention on these two types of responses. As suggested by the differential responsiveness analysis (Figure 4c), we found that most significant responses to visual and standard-1 stimuli were observed only in one of the two states—not both (Figure 4i).

We then examined whether the visual and standard-1 responses were localized to specific thalamic nuclei. We limited our analysis to seven regions (AV, CL, LD, LHB, MD, VL and VPM) for which we had sufficient data from at least eight recordings from at least four mice. Surprisingly, visual responses were found widespread across recorded thalamic nuclei: all nuclei responded strongly under anesthesia but much less so during wakefulness (Figure 4j). Significant auditory responses were observed in AV, CL, LHB, MD, and VPM in awake state, and anesthesia reduced the responses in all these areas but CL (Figure 4j).

In sum, thalamic nuclei respond in a stimulus- and state-dependent manner; anesthesia generally reduces auditory responses but enhances visual responses.

### Locomotion modulates multi-unit activity across thalamic nuclei

When awake, animals were free to move on a treadmill at will, whilst we recorded the movement of the treadmill together with LFP and MUA in the thalamus. To examine how movements (and accompanying sensory inputs) affect the activity across thalamic nuclei, we compared the average MUA during moving and stationary periods for every recording site (Fig 5a). We found significantly higher MUA during periods of movement across many thalamic nuclei (Fig 5a). Specifically, we found that 194 out of 352 sessions showed significantly movement-modulated activity with most sessions had higher MUA with some exceptions that showed a decrease (two-sided Mann-Whitney U test, n=352 sites across 11 mice); even with higher p-values, a large fraction of movement-responsive electrode sessions remained movement responsive (Fig 5b).

**Figure 5:**
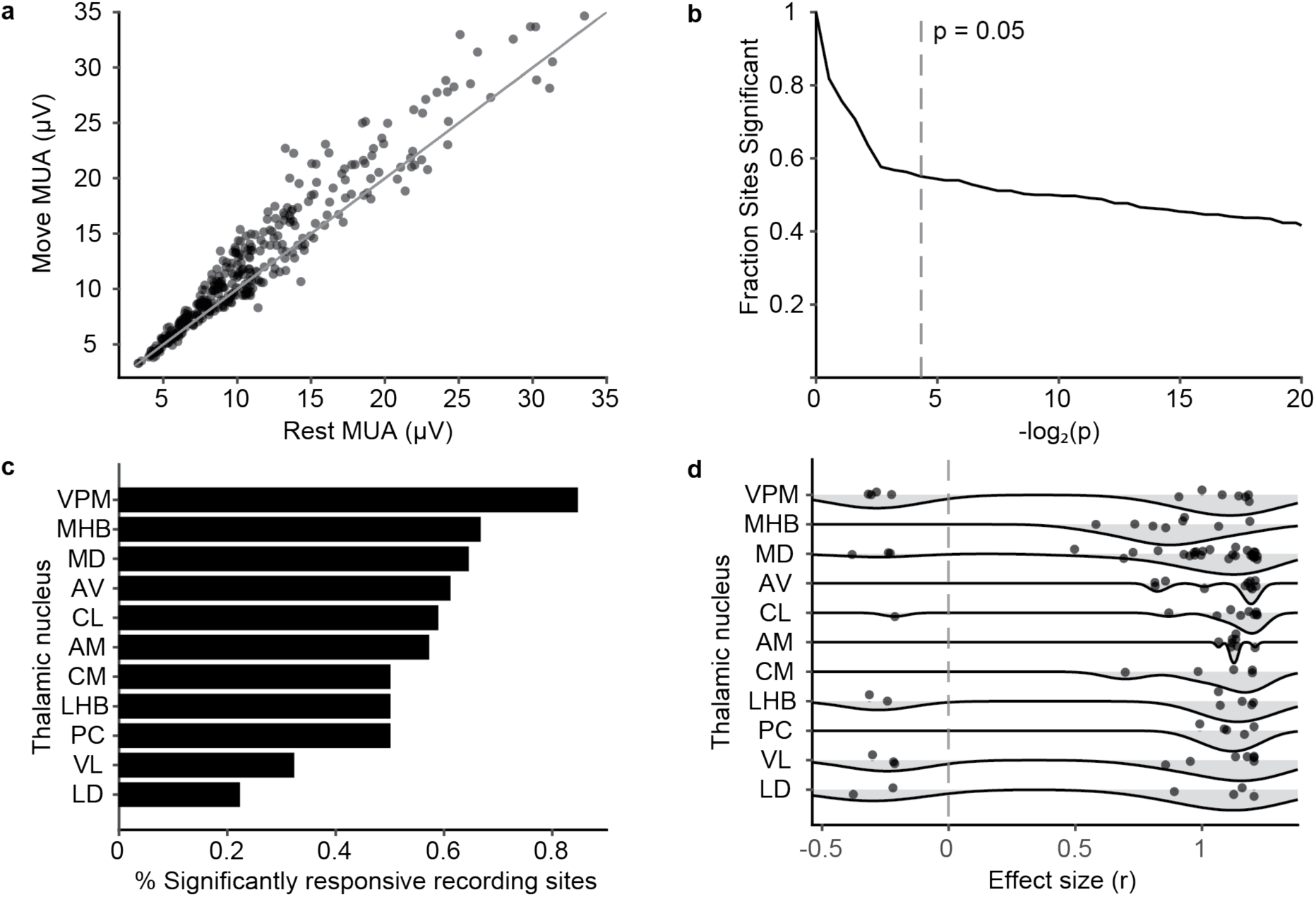
Locomotion enhances MUA across thalamic nuclei. (a) Average MUA during periods of movement (see methods) and periods of rest (n=352 sites, n=11 mice). Every dot represents one recording site. (b) Fraction of recording sites showing significant activity related to movement across different significance thresholds (n=352, two-sided Mann-Whitney U test, FDR corrected with the Benjamin-Hochberg method). The dashed line indicates α=0.05. (c) Movement responsiveness across thalamic nuclei. Shown are fractions of significantly responsive (α=0.05) sessions in different thalamic nuclei, with the highest fraction in the VPM but also movement related responses in other thalamic nuclei. (d) Distribution of effect sizes (see Methods) for recording sites in different thalamic nuclei (same as in C) in response to movement. Every dot represents one recording site in each thalamic nucleus. Nuclei mostly increase their activity during movement, but notably some sites also show significantly decreased activity.

Subsequently, we wondered if these movement responses were specific to certain thalamic nuclei only. We examined the thalamic nuclei whose data were from at least 4 animals and at least 10 total recordings. As expected, the first-order somatosensory nucleus VPM contained the highest fraction of movement-responsive sites (Fig 5c), but interestingly movement modulated all recorded thalamic nuclei, including those that are not classically described as motor nuclei—such as MD, CL, and CM (Fig 5c). All nuclei showed higher MUA during locomotion, but 45% (5/11) of nuclei as well as LHB also had neurons of decreased MUA during locomotion (Fig 5d).

### Modality- and state-specificity of thalamic responses

Finally, we examined the specificity of neuronal response in each thalamic nucleus. So far, we observed three main drivers of MUA responses: the visual flash under anesthesia (57% of differentially responsive sites), the standard-1 tone during wakefulness (42%) and movement on the treadmill (61%, assessment possible during wakefulness only). However, some thalamic nuclei may selectively respond to a particular stimulus type in one conscious state whereas other nuclei may respond more broadly in other conditions. Since we were surprised by the large fraction of recording sites responding to the visual flash under anesthesia, we were especially interested in the response patterns of those nuclei. We based this analysis on our responsiveness analysis from Fig 4 which included data from 191 recording sessions with significant stimulus responses in at least one state.

An overview of the responsiveness for each recording site to each condition is shown in Fig 6a. We calculated the pairwise frequencies of recording sites responding to two conditions at the same time (Fig 6b) and arranged them into a graph to look for functional clustering (Fig 6c). We observed a cluster of recording sessions (n=27) that responded to the three main drivers (the visual flash during anesthesia, the standard-1 tone during wakefulness and movement on the treadmill) (Fig 6d).

**Figure 6:**
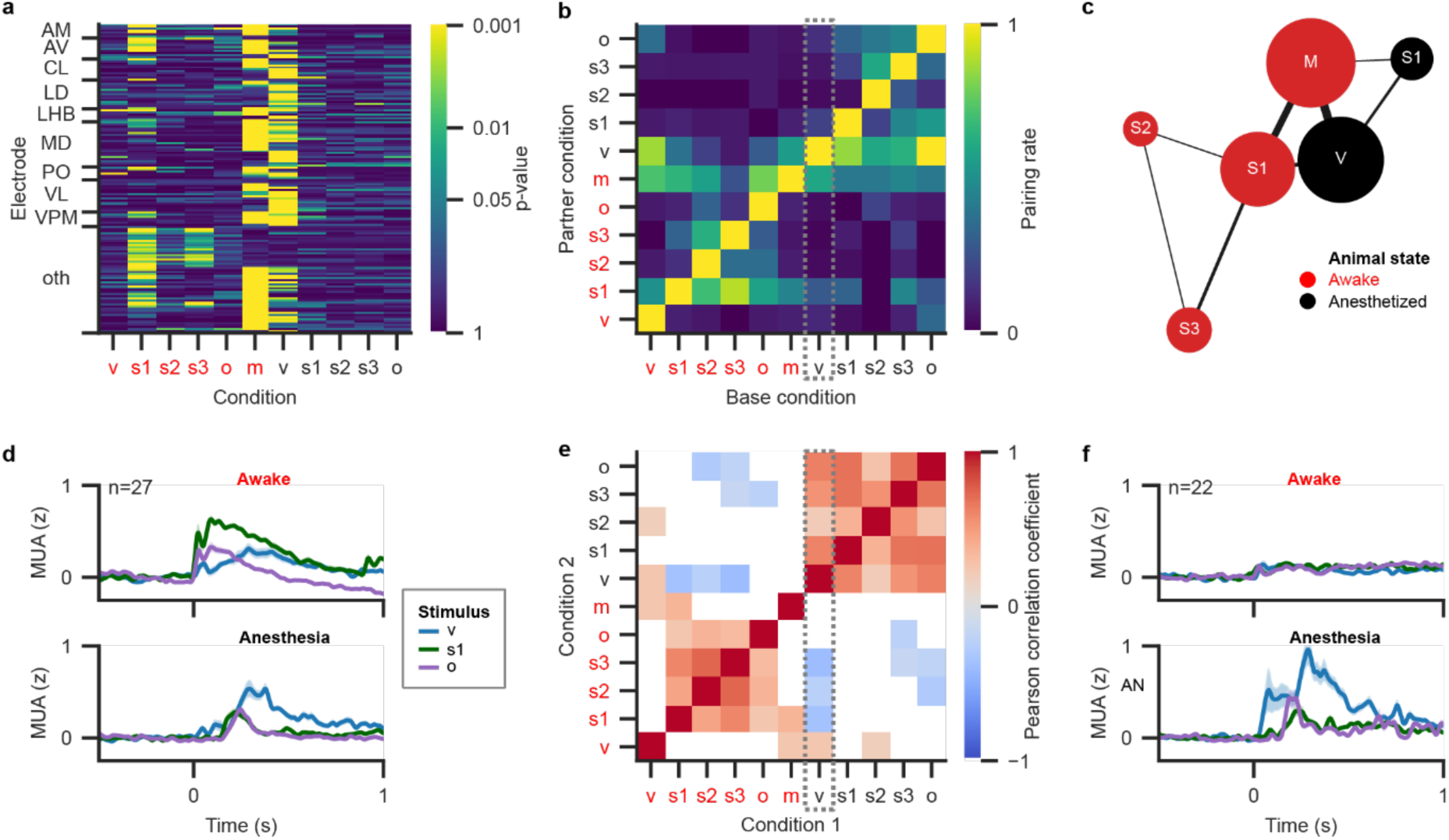
The selectivity of thalamic neurons. (a) p-values of 191 significantly responsive sites that indicate responsiveness eleven conditions. p-values of stimulus conditions were calculated by Mann-Whitney U test between MUA responses in null and stimulus trials (see also Fig 4 & Methods). p-values for the treadmill condition were calculated by Mann-Whitney U test between MUA responses during periods of movement and no movement on the treadmill (see also Fig 5 & Methods). All p-values were corrected for multiple comparisons (see Methods). Red: wakefulness, black: anesthesia, v: visual flash, s1: standard-1 tone, s2: standard-2 tone, s3: standard-3 tone, o: oddball tone, m: movement on treadmill. Electrodes are sorted hierarchically by nucleus and condition of smallest p-value. oth: recording sites in nuclei with fewer than eight sites or missing area information. (b) Pairing rate of recording sites responding to pairs of conditions, where a value of 1 indicates complete overlap in responsiveness and 0 indicates no overlap. Abbreviations as in a. Dotted rectangle highlights the column with pairing to responsiveness to visual flash under anesthesia. M = electrodes responsive to movement on the treadmill (n = 116), S1 = electrodes responsive to the standard-1 tone (awake: n = 80, anesthetized: n = 25), S2 = electrodes responsive to the standard-2 tone (awake: n = 16), S3 = electrodes responsive to the standard-3 tone (awake: n = 28), V = electrodes responsive to the visual flash (anesthetized: n = 108). (c) Graph of shared responsiveness of recording sites. Nodes represent conditions. Node size indicates the number of sites responsive to that condition. Edge width indicates the number of shared sites between two conditions. For visualization purposes, only conditions with at least 15 responsive recording sessions and connections with at least 10 shared electrodes are shown. Electrodes often responded to all three main drivers of MUA (visual flash during anesthesia, standard-1 tone and treadmill movement during wakefulness). (d) Average MUA responses of 27 recording sites that significantly respond to the three main drivers of MUA. AW: wakefulness, AN: anesthesia. Other abbreviations as in (a). The responses were smoothed with a 30-ms Gaussian kernel before averaging for visualization purposes. (e) Pearson partial correlation coefficients of response effect sizes between condition pairs. Only significant coefficients are shown, all coefficients with pcorr > 0.05 were set to white for visualization purposes. The dotted rectangle highlights the column of correlations of the responsiveness to the visual flash stimulus under anesthesia. r = Pearson partial correlation coefficient. (f) Same as (d) but now for 22 recording sites that only responded significantly to the visual flash under anesthesia.

We wondered whether this triple association meant that recording sites preferentially responded to those conditions together. We therefore measured partial correlations between responses to all condition pairs while correcting for animal specific effects (Fig 6e, see also Methods). Surprisingly, despite often co-occurring, neither responsiveness to the standard-1 tone nor responsiveness to treadmill movement during wakefulness correlated positively with responsiveness to the visual flash under anesthesia. In contrast, responsiveness to the visual flash under anesthesia was positively correlated with responsiveness to all other anesthetized conditions, highlighting again that stimulus signals were processed in a state-dependent manner in the thalamus. Consistently, we found recording sessions with significant responses only to the visual flash under anesthesia and to no other condition (Fig 6f). These results suggest that the subpopulation of thalamic neurons that respond during anesthesia are different from the subpopulation that respond in awake states.

## Discussion

Our findings challenge the classical view of the thalamus as a collection of independent nuclei^18^ and rather suggest instead that it operates as a coordinated structure which is surprising in light of the lack of local excitatory recurrent connections within the thalamus as well as the sparsity of GABAergic neurons^18^ (further discussion on the potential source of coordination will follow). We observed strikingly correlated LFP across multiple thalamic nuclei, which were strongly influenced by behavioral state and sensory input. This finding is supported by several observations: significant correlations in LFP and MUA across nuclei, extensive MUA responses to movement and sensory stimuli, and prominent state- and modality-dependent responses. Together, these results suggest coordination of activity patterns across different thalamic nuclei.

Many thalamic nuclei showed a significant decrease in MUA activity during anesthesia. This widespread reduction is consistent with previous studies of individual HO thalamic nuclei associated with consciousness^10–17,22^, implying their common role in maintaining conscious states. This idea is further supported by our observation of strong correlations in LFP across anatomically distinct nuclei, suggesting the existence of shared synaptic inputs to different thalamic nuclei. Volume conduction could partly explain the correlation^23–27^, which could originate from a source in the thalamus itself as well as in cortical areas, although it is unlikely to come from within the thalamus due to the anatomical ineffectiveness of the thalamus to evoke LFP^28^.

However, the highly correlated LFP might be caused by strong distant sources outside the thalamus such as the hippocampus or cortical areas^25^. Although volume conduction can partly contribute, all the analyses we conducted suggest the existence of local (within-thalamus) nature of this signal for three reasons. First, both monopolar and bipolar LFP are significantly correlated even between long distant (up to 1,600μm) thalamic nuclei. Second, comparing the correlations between bipolar LFP electrodes in the thalamus and a control shank in the cortex suggests that this correlation was thalamus specific. Even after removing the most prominent global signal (calculated by ICA and other methods) across the cortex and the thalamus, the intra-thalamic correlations remained significantly higher than the thalamo-cortical correlations. Lastly, we found a significant correlation of MUA across different thalamic nuclei. This residual, thalamus-specific correlation as well as the MUA correlation between thalamic nuclei could thus suggest the presence of shared inputs within the thalamus.

Additionally, widespread MUA responses to locomotion and sensory stimuli were observed across a variety of thalamic nuclei, including those that have traditionally not been associated with motor or specific sensory functions. Such activity patterns across nuclei are more consistent with a coordinated network rather than independent, parallel channels. Notably, state changes dramatically altered the sensory modality that thalamic nuclei responded to: under anesthesia, a wide range of thalamic nuclei responded robustly to visual stimuli, whereas auditory responses were mostly present during wakefulness. Moreover, the most thalamic nuclei responded only in either anesthesia or wakefulness, which suggests that distinct neuronal populations, present across nuclei, selectively responded to different stimuli. The nature of these shared responses appears complex and context-dependent. For instance, the widespread response to auditory stimuli may partly arise from associated motor activity, as sounds frequently trigger stereotyped movements^29–32^. Indeed, substantial overlap was observed between channels responsive to auditory stimuli and those active during locomotion. Whilst responsiveness to movement was positively correlated with responsiveness to the first auditory stimulus, suggesting a possible dependence between the two responses, the responses to the visual flash during anesthesia was not correlated with movement nor auditory responses during wakefulness, suggesting the independence of this response.

Which mechanisms could explain this apparent coordination across thalamic nuclei? The correlated LFPs could suggest the regulation of different thalamic nuclei by common sources. Individual thalamic nuclei receive diverse nucleus-specific inputs from cortical and subcortical regions^33^, but in addition to nucleus-specific inputs, thalamic nuclei receive common neuromodulatory inputs^34^. Moreover, all thalamic nuclei receive inhibitory projections from caudal prethalamic regions, especially the thalamic reticular nucleus (TRN) and zona incerta^35^. The TRN has been proposed to regulate information flow from cortex to thalamus^36^, potentially synchronizing rhythms across nuclei despite nucleus-specific inputs. Neuromodulatory systems may similarly coordinate activity of caudal prethalamic regions. Importantly these caudal prethalamic regions also project to the habenula^35^, which could possibly explain why this region showed the similar state-dependent responses. Caudal prethalamic regions (such as the TRN and zona incerta) have been described as inhibitory switchboards, modulating their target output based on behavioral demands^35^. These regions could adjust activity patterns depending on the animal’s state, thus driving the observed state-dependent responses. Indeed, the TRN itself exhibits state-dependent shifts, altering its inhibitory targeting of different nuclei during sleep versus wakefulness^37^. An anesthetic-induced state shift, possibly due to the changes in neuromodulatory input, in TRN or other caudal prethalamic nuclei may represent a plausible mechanism underlying the state-dependent thalamic responses we observed.

The anatomical fact that different thalamic nuclei project to diverse cortical regions^38^ implies that their activity patterns likely induce widespread modulations across the cortex. A previous hypothesis posits that the thalamus facilitates belief updating in the cortex^39^. Within this framework, activation of multiple thalamic nuclei by sensory stimuli and movement may reflect the brain’s widespread need for arousal-related signals to dynamically adjust cortical processing. In fact, it has been shown that arousal reduces perceptual bias^40^, suggesting that arousal can alter ongoing sensory processing. Moreover, HO thalamic nuclei have already been linked to arousal^10,11,17,41,42,43^. For example, L6 pyramidal neurons in S1 send arousal-related information to one of HO nuclei (POm), which in turn can increase cortical activity^44^. This suggests that HO nuclei could integrate both sensory and behavioral state information to regulate cortical activity in a context-dependent manner.

In summary, our findings challenge the prevailing view of strictly independent functioning of higher-order thalamic nuclei. We found that multiple HO nuclei simultaneously responded to sensory stimuli and correlated LFPs between these different nuclei, suggesting the existence of a mechanism that coordinates thalamic nuclei. Future studies may address the mechanism that coordinates the HO nuclei and their downstream cortical areas.

## Methods

### Animal Models

All animal experiments were performed according to the national and institutional regulations. The experimental protocol was approved by the Dutch Commission for Animal Experiments and by the Animal Welfare Body of the University of Amsterdam. Adult wild-type female and male mice were used in this study.

### Extracellular recordings

A four-shank electrode with eight channels per shank (A4x8-5mm-200-400-177-A32, NeuroNexus) was targeted to the higher-order thalamic nuclei using a micromanipulator (SM-15M, Narishige, Tokyo). The extracellular potentials were digitized and amplified by a RHD2132 and RHD USB interface board.

### Drug application and sensory stimulation

General anesthesia was induced by isoflurane (induction at 3%, maintenance at 1.5-2%). Visual stimulation was applied by an LED-coupled optical fiber pointing at the right eye of the mouse (distance: 1-2 cm). The auditory stimulus was presented using an 8 Ohm speaker placed ∼30 cm away from the mouse.

### Sensory stimuli and behavior

Mice were presented with visual and auditory stimuli. The visual stimulus was a 100 ms flash of white light. The auditory stimulation consisted of an oddball paradigm with a random number (between 3 and 5) of tones at 2 KHz followed by one tone at 4 KHz; all tones lasted for 100 ms, and the inter-tone interval was 900 ms. The inter-trial interval was randomly chosen from 3–5 seconds. Each session consisted of 10 auditory and 10 visual trials and at least 3 sessions were recorded from each mouse in both anesthesia and wakefulness. Moreover, awake mice could freely walk or run on a linear treadmill at any point during the recording.

### Verification of electrode location

After each recording, the probe was coated with DiI (Merck) and inserted to verify the locations of the electrodes. Brains were cut into 100 micrometer coronal sections. Subsequently we overlaid the photograph of the slice with the Paxinos and Franklin mouse brain atlas (2001). We then fitted the diagram of the 4-shank probe on the DiI traces to determine the position of the 4-shank electrodes.

### Data Pre-processing

We used MATLAB R2024B (Mathworks, 2024) for data pre-processing. Data were first separated into LFP and MUA. The LFP was extracted by low-pass filtering (<200Hz) the raw signal using a fifth-order Butterworth filter in MATLAB. The MUA was extracted by bandpass filtering the signal between 300 and 5000 Hz using a third-order Butterworth filter, after which it was re-referenced to the common-average of the shank to reduce artefacts. Lastly, the signal was full-wave rectified and low-pass filtered at 200 Hz with a third-order Butterworth filter to obtain the MUA envelope. This MUA envelope reflects the average spiking activity of neurons in the vicinity of the recording site^45^.

### Spontaneous activity analysis

Firstly, we averaged MUA across electrodes and compared this between wakefulness and anesthesia using a two-sided Wilcoxon signed-rank test; the same approach was used using the root-mean-square LFP. Subsequently, we calculated the power spectrum in both states using Welch’s method, after which we calculated the Delta Ratio as the activity in the Delta range (0.5 -4 Hz) divided by the power in the rest of the sub-30 Hz spectrum. To determine the effect of anesthesia on the delta ratio, we used a linear mixed effects model with state (anesthesia/awake) as the predictor (independent variable), the delta ratio as the predicted (dependent) variable and the animal ID as a random effect to account for differences between animals.

### Correlation analysis

Correlation coefficients were calculated per recording session between all sites and then averaged over sessions. We used Pearson correlation as our method to quantify correlations. For the frequency band analysis, we first filtered the signal in the different frequency ranges using a fourth-order bandpass Butterworth filter, where delta was defined as <4Hz, Theta 4-8 Hz, Alpha 8-12 Hz, Beta 12-30 Hz, Gamma 30-80 Hz. Analyses were performed on monopolar re-referenced LFP where the reference electrode was placed above the cerebellum, bipolar re-referencing was applied by subsequently subtracting signals from adjacent electrode contacts along each shank, yielding 28 pairs for 32-channel thalamic arrays and 43 pairs for 48-channel thalamic + cortical arrays.

To further control for volume-conduction, an additional electrode shank was placed in the cortex (n=3 mice). For this cortex-control we used the same approach of calculating the correlation between all electrode pairs. Afterwards, we differentiated between thalamus-thalamus pairs, thalamo-cortical pairs and intra-cortical pairs to compare intra-thalamus correlations with thalamo-cortical correlations, to assess to what extent the LFP correlations were specific to thalamic recording sites.

For the population coupling analysis, we first created the population signal by summing the MUA signals across all (except for the cortical control) electrodes. Then, we correlated the signal from every electrode with the population signal using the Pearson correlation which granted their population coupling.

### Correlation over distance

The relation of signal correlation with anatomical distance was determined using a linear mixed effects model with correlation in LFP between electrodes as the dependent variable, the distance between electrodes (vertical or absolute) as the predictor (independent variable) and the animal identity as a random effect.

### Analysis of MUA stimulus responses

MUA stimulus responses were analyzed using Python 3. For each trial, the MUA was z-scored by using the mean and standard deviation of 500 ms pre-stimulus onset (500 – 0 ms before stimulus onset). Subsequently, we created a null condition from the pre-stimulus, pre-scaling period of each visual trial (1000 – 500 ms before stimulus onset).

To assess whether a recording site showed significant stimulus responses, we calculated a Kruskal-Wallis test for each recording site and state separately. In each trial, we selected the z-scored MUA from 0 to 500 ms post stimulus onset. To avoid having negative and positive z-values cancel out, we squared the MUA values and then averaged them for each trial. This way, we ended up with a single MUA value for each trial. We then grouped the MUA values by stimulus type, which resulted in six groups (five stimulus conditions: visual flash, standard-1 tone, standard-2 tone, standard-3 tone, oddball tone; one null condition). Each group had at least thirty trials. We computed the Kruskal-Wallis test over the six groups. We then corrected the p-values from all tests (344 electrodes x 2 states = 688 test) using Benjamini-Hochberg FDR correction. An electrode was considered as responsive in a state if its corrected Kruskal-Wallis test p-value was smaller than 0.05.

Next, for all 191 electrodes that were found to be responsive in at least one state, we computed pairwise two-sided Mann-Whitney U tests between stimulus and null responses. We again took the average squared z-scored MUA from 0 to 500 ms post stimulus onset in each trial. For each electrode, stimulus type and state, we computed a two-sided Mann-Whitney U test between the MUA responses to that stimulus and the MUA responses in the null condition. Additionally, we also computed the effective difference between the MUA responses of the stimulus and null conditions as quantified by Cohen’s d. For each electrode in each state, we ended up with five p-values (one for each stimulus type). We corrected these five p-values using Benjamini-Hochberg FDR correction. An electrode was considered as significantly responsive to a stimulus type in a given state if its corrected Mann-Whitney U test p-value was smaller than 0.05.

### Movement responses

Movement was classified from the treadmill signal by first filtering the data to highlight movement related bursts, then computing a smoothed amplitude envelope. We standardized this envelope using a robust z-score, and applied hysteresis thresholds to decide when movement started and stopped. To improve reliability, very short bursts were discarded, closely spaced bursts were merged, and regions near the signal edges were ignored. The result is a clean binary classification of movement versus non-movement. Movement responsiveness was determined for each electrode by computing the mean MUA activity in periods of 1,000 ms with an equal amount of non-movement periods and then comparing movement- and non-movement MUA using two-sided Mann-Whitney U tests. We then corrected p-values using Benjamini-Hochberg FDR correction, with an alpha of 0.05. The effect size was quantified as 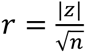, where z is the Mann-Whitney U Test z-value.

### Partial correlations

In “*Analysis of MUA stimulus responses*”, we explained how we computed the effective difference (quantified as Cohen’s d) between stimulus and null MUA responses for each electrode, stimulus type and state separately. For the analysis of partial correlations, we took this effective difference as a continuous measure of responsiveness to that condition. There were 11 conditions in total: 5 stimulus types x 2 states + 1 treadmill condition. To assess whether responsiveness in one condition was correlated with responsiveness in another condition, we computed the Pearson partial correlation coefficient (and its associated p-value) between the effective difference vectors of every condition pair, taking the animal ID as a co-variate to correct for animal-specific effects. Each effective difference vector contained 191 elements, one for each differentially responsive electrode. There were 55 unique condition pairs, and we corrected the p-values of those pairs using Benjamini-Hochberg FDR correction.

## Author contributions

T.B: Data curation, Formal analysis, Visualization, Writing – original draft

M.B: Data curation, Formal analysis, Visualization, Writing –results section, review & editing

C.M.A.P: Funding acquisition, Writing – review & editing

M.S: Conceptualization, Investigation, Supervision, Funding acquisition, Writing – review & editing

## Acknowledgments

M.S. discloses support for the research of this work from Research Foundation for Opto-Science and Technology, Kayamori Foundation of Informational Science Advancement, Mitsubishi Foundation, Dutch Research Council, Brain and Behavior Research Foundation, International Brain Research Organization, Japan Society for The Promotion of Science, Takeda Science Foundation, Naito Foundation, Sekisui Chemical, Daiichi Sankyo Foundation of Life Science, and Nakajima Foundation.

## Supplementary figures

**Supplementary figure 1:**
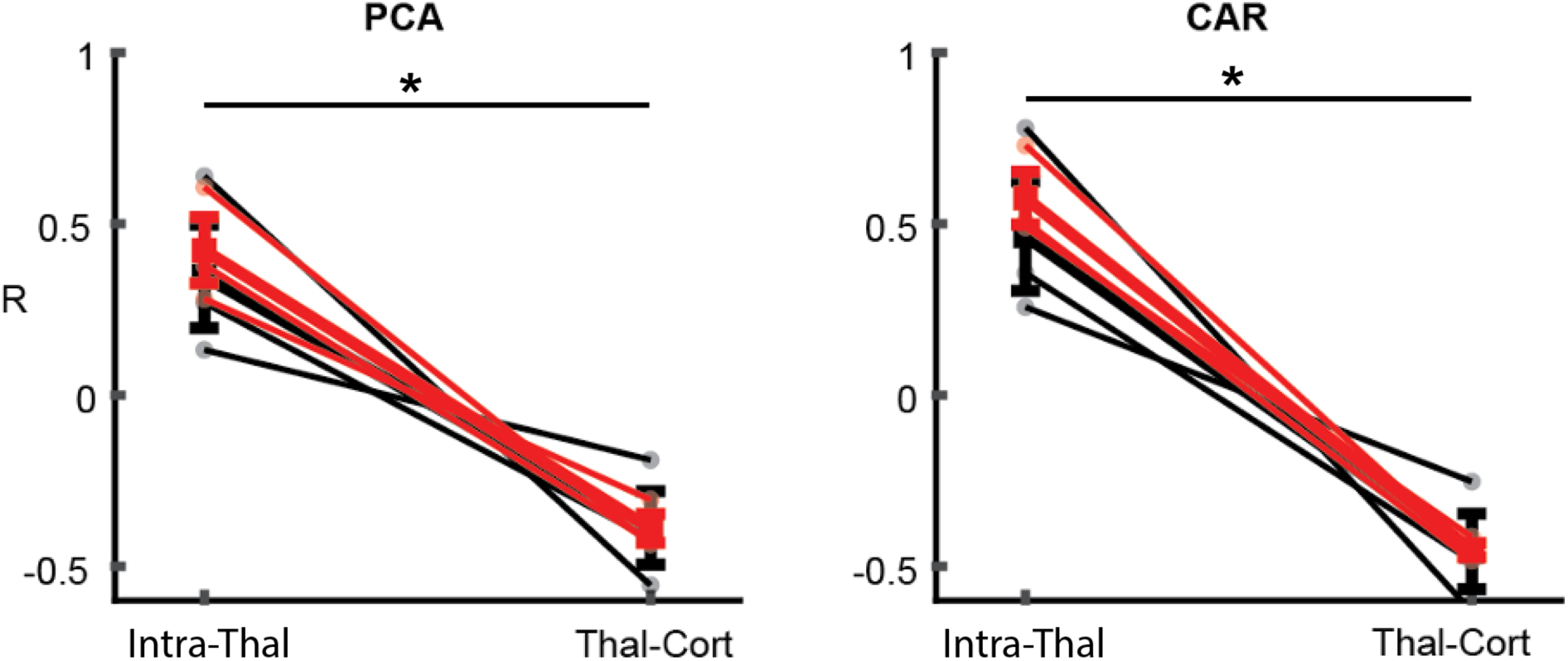
Alternative methods for examining volume conduction. Left: The intra-thalamic correlation (R) remained high even after removing the first principal component extracted from the LFP data from all recording electrodes in both cortex and thalamus. Right: Same as in the left but now the common average across all electrodes in both thalamus and cortex is removed. CAR, common average re-referencing (see Methods).

